# Functional recovery associated with dendrite regeneration in PVD neuron of *C. elegans*

**DOI:** 10.1101/2023.08.09.552579

**Authors:** Harjot Kaur Brar, Swagata Dey, Pallavi Singh, Devashish Pande, Anindya Ghosh-Roy

**Affiliations:** Department of Cellular & Molecular Neuroscience, National Brain Research Centre, Manesar, Haryana-122052, India; Current address: Department of Psychological and Brain Sciences, University of Delaware, Newark, Delaware- 19716, USA

**Keywords:** *C. elegans*, PVD neuron, dendrotomy, dendrite regeneration, harsh touch, posture

## Abstract

PVD neuron of *C. elegans* is a highly polarized cell with well-defined axonal, and dendritic compartments. PVD neuron operates in multiple sensory modalities controlling both nociceptive touch sensation and body posture. Although both axon and dendrites of this neuron show regeneration response following laser-assisted injury, it is rather unclear how the behavior associated with this neuron is affected by the loss of these structures. It is also unclear whether neurite regrowth would lead to functional restoration in these neurons. Upon axotomy, using a femtosecond laser, we saw that harsh touch response was specifically affected leaving the body posture unperturbed. Subsequently, recovery in the touch response is highly correlated to the axon regrowth, which was dependent on DLK-1 MAP Kinase. Dendrotomy of both major and minor primary dendrites affected the wavelength and amplitude of sinusoidal movement without any apparent effect on harsh touch response. We further correlated the recovery in posture behavior to the type of dendrite regeneration events. We found that dendrite regeneration with the fusion and reconnection between the proximal and distal branches of the injured dendrite corresponded to improved recovery of posture. Our data revealed that the axons and dendrites differentially regulate the functions of PVD neurons. It also revealed that dendrite and axon regeneration are both functionally and molecularly distinct.

## Introduction

Both axons and dendrites are vulnerable to damage upon accidental injury (Mizielinska et al., 2009, Park et al., 1996, Risher et al., 2010). The molecular mechanism of axon regeneration is highly studied over last few decades as behavioral restoration after neuronal injuries highly depends on the regrowth from the injured axonal stump and its rewiring (Laha et al., 2017, Rasmussen and Sagasti, 2017, Becker et al., 1997, Yanik et al., 2004, Basu et al., 2021) (Zheng and Tuszynski, 2023, Winter et al., 2022). Manipulation of the cAMP pathway, mTOR signaling, and epigenetic molecules led to functional recovery following the injuries to central nervous system in vertebrate models (Weng et al., 2017, Bei et al., 2016, Qiu et al., 2002). The use of finely focused lasers like a 2-photon microscope in the precise injury of axons in models such as nematode, fruit fly, and zebrafish opened the window to understand the basic biology of axon regeneration (He and Jin, 2016, Ghosh-Roy and Chisholm, 2010, Richardson and Shen, 2019, Yanik et al., 2004). A p38 MAP kinase cascade involving Dual leucine zipper kinase kinase kinase (DLK-1) is essential for initiating axon regeneration from the cut tip of the axon in various models (Hammarlund et al., 2009, Yan et al., 2009, Ghosh-Roy and Chisholm, 2010, Ghosh-Roy et al., 2010, Shin et al., 2012). Axon regeneration potential declines with age at both anatomical and functional levels (Verdu et al., 1995, Geoffroy et al., 2016, Basu et al., 2021). Manipulation *let-7* miRNA, Insulin (IIS) signaling and AMP kinase signaling can help overcome the age-related decline in axon regeneration potential in *C. elegans* (Zou et al., 2013, Basu et al., 2017, Wang et al., 2018, Kumar et al., 2021). In comparison to axon regeneration, the regeneration response following a physical injury of dendrites are less studied until recently (Thompson-Peer et al., 2016, Oren-Suissa et al., 2017, Stone et al., 2022).

The highly branched dendrites of Drosophila sensory da9 neurons and dendrites of PVD neurons in *C. elegans* can regrow following laser-assisted injury (Thompson-Peer et al., 2016, Oren-Suissa et al., 2017, Brar et al., 2022). Both in fly and worms, dendrite regeneration is independent of the conserved DLK-1 MAP Kinase pathway (Stone et al., 2014, Brar et al., 2022), which is essential for axon regeneration in multiple organisms (He and Jin, 2016). Dendrites of motor neurons can regenerate in the zebrafish spinal cord (Stone et al., 2022). Therefore, understanding the molecular mechanism regulating dendrite regeneration and its functional significance are endeavors of high importance. After the dendrotomy of class IV da neuron in the fly, the nociceptive function is affected, which subsequently is recovered through dendrite regrowth (Thompson-Peer et al., 2016, Hertzler et al., 2023). However, these experiments are done in larval stages. Therefore, regrowth seen in fly dendrites after laser injury could partly be overlapping with the remodeling process during larval development (Yaniv and Schuldiner, 2016).

PVD neurons in *C. elegans* are recognized for a range of behavioral responses to external stimuli, including harsh touch response, thermo-sensation, sound sensing, and proprioception (Chatzigeorgiou et al., 2010, Iliff et al., 2021, Albeg et al., 2011, Tao et al., 2019, Ibsen et al., 2015). When a worm is subjected to a harsh touch using a platinum wire, it exhibits an escape response behavior. PVD plays an important role in sensing the harsh touch (Li et al., 2011, Tao et al., 2019). The TRP-4 (DEG/ENAC) channel in PVD is responsible for detecting the harsh touch, resulting in the inflow of Na2+ and K+ ions (Li et al., 2011). Another study indicated that DEG/ENaC channel, ASIC-1, and the TRPM channel, GTL-1 play an important role in shaping excitability in response to touch (Husson et al., 2012).

PVD neurons also regulate body posture during movement (Albeg et al., 2011, Tao et al., 2019). The sinusoidal waves generated by a moving worm are the signature of the proprioception behavior (Albeg et al., 2011). PVD neuron is responsible for the optimal values of the amplitude and wavelength of the sinusoidal traces made by a moving worm on a bacterial lawn (Albeg et al., 2011, Tao et al., 2019). According to a recent analysis, dendrites regulate neuromuscular junctions, which govern the excitation and inhibition of body wall muscles (Tao et al., 2019). In response to the worm’s sinusoidal movement, the PVD dendrites generate local calcium influx and release neuropeptide, NLP-12. NLP-12 neuropeptide directly modulates neuromuscular junction activity through the cholecystokinin receptor on motor axons. (Tao et al., 2019). This local calcium influx is dependent on the DEG/ENaC channels MEC-10, UNC-8, and DEL-1 (Tao et al., 2019). The same study indicated that the DEG/ENaC/ASIC channel, DEGT-1 acts as a mechanoreceptor for harsh touch, which is transduced through the axon to the downstream interneuron (Tao et al., 2019). Therefore, the dendrites and axons of PVD neurons could differentially regulate the proprioception and harsh touch sensation behaviors independently.

In this work, we wanted to get insight into how axotomy and dendrotomy would impact PVD-related behaviors. We analyzed both harsh touch sensation and body posture after the axotomy or dendrotomy using a femtosecond laser. We discovered that laser injury to primary dendrites exclusively affects body posture while leaving the harsh touch response unchanged. Whereas the axotomy affects harsh touch sensitivity without altering the proprioception. At later time points, the animals suffered from axotomy-induced loss of touch sensation regained their function in a DLK-1/MLK-1 MAP Kinase-dependent manner. The behavioral recovery related to proprioception after dendrotomy depends on the pattern of the regeneration response. The recovery in body posture parameters is highly correlated with the fusion events between the proximal and distal dendritic branches of the injured PVD. Our data highlights the functional significance of dendrite regeneration in adulthood.

## Results

### Harsh touch response is affected by axotomy of PVD neuron but it remains unaffected by dendrotomy

PVD neuron displays an orthogonal array of dendritic branches that cover major part of the body (Fig 1A) (Inberg et al., 2019). The higher order branches are arranged in a menorah like fashion (Fig 1A) (Oren-Suissa et al., 2010). Since both harsh touch sensation and proprioception are largely controlled by PVD neurons it is not clear whether one or both the modalities will be affected by dendrotomy or axotomy. Therefore, to understand whether dendrite regeneration would lead to functional restoration, it is important to have a quantitative behavioral deficit in a specific behavior caused due to dendrotomy. We first studied the effect of axonal and dendritic injury on the harsh touch response behavior. When the worms were prodded near the vulva with a platinum wire, they instantly exhibited an escape response (Fig. 1B, red mark, Movie S1) as seen before (Li et al., 2011). The distance traveled by the worm in each frame was shorter before the delivery of the harsh touch stimulus but it increased thereafter (Fig. 1C, Movie S1). Nearly 90% of the wildtype worms responded to the harsh touch (Fig. 1D) and the response remained comparable in the L4 and Day-1 stages, with a 20% decrease at the Day-2 stage (Fig. 1D). The percentage of animals responding to harsh touch was significantly reduced in *mec-3* mutant (Fig. 1D). The *mec-3* gene codes for a transcription factor, which is required for the expression of genes important for the development and function of PVD (Smith et al., 2013). However, this sensory behavior was unaffected in the gentle touch defective mutant of *mec-4* (Fig. 1D), which codes for the DEG/ENaC mechanoreceptor channel for gentle touch (Chalfie and Sulston, 1981). This indicated that our assay condition is specifically sensitive to harsh touch.

**Figure 1:**
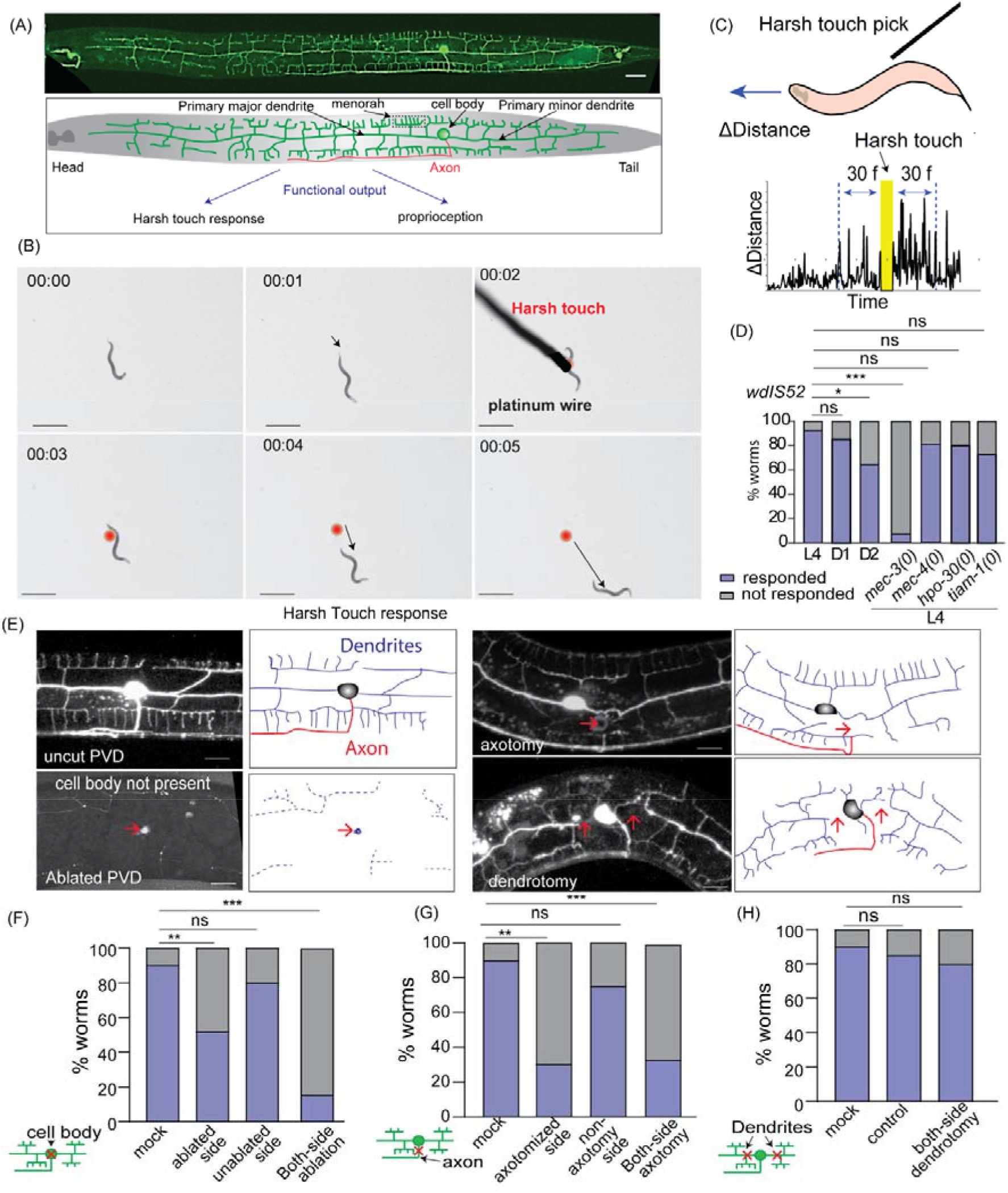
Nociceptive response to harsh touch depends on the integrity of PVD axons but not the dendrites. (A) Confocal image of PVD neuron expressing *wdIs52* (*pF49H12.4:: GFP*) reporter and the schematics showing dendrites in green, and axon in red color. The scale bar represents the 20 microns value. (B) Time-lapse video montage showing worm’s response before and after the harsh touch by platinum wire is shown. The worm is placed on the NGM plate. The black arrow represents the worm’s distance covered in each frame. The red dot indicates the location of the worm where the harsh touch was made. The scale bar is 1 mm. Time-frames are indicated in min:sec. (C) Schematics representing the harsh touch pick and instantaneous distance travelled after each frame. The representative plot in the bottom shows the distance travelled by the worm in each frame vs time. The yellow bar shows the harsh touch event. The blue dotted lines represent the 30 frames duration taken to calculate the speed of the worm. (D) The percentage of worms responding to the harsh touch was plotted on contingency-based graphs for L4, Day1 adult, Day2 adult worms along with *mec-3(0)*, *mec-4(0)*, *hpo-30(0),* and *tiam-1(0)* at L4 stage, 15<n<25, N=3. (E) The confocal images of uncut, ablated, axotomized, and dendrotomized PVD neuron at 3-6h after injury is shown along with the illustrations for dendrites and axon. The red arrow marks the site of injury. The scale bar represents 20 microns. (F-G) Percentage of L4 *wdIs52* transgenic worms responding to harsh touch with one or both PVD neurons ablated (F) or axotomized (G) compared to mock injured worms. For one sided injured worms, harsh touch was given on both injured (ablated or axotomized) and uninjured (unablated or non-axotomy) side. The data is plotted in a population-based contingency plot. 15<n<25, N=3. The type of injury is shown in schematics with the red cross mark. (H) Percentage of L4 *wdIs52* worms responding to harsh touch in control (uninjured), mock, and both-side (PVDL and PVDR) dendrotomy injury conditions. 12<n<28, N=3. The type of injury is shown in schematics with the red cross mark. The data is plotted in a population-based contingency plot. The statistical analysis for D, F, G, and H, was Fisher’s two-tailed exact test with p value as p < 0.05*, 0.01**, and 0.001***. ns = not significant, N = independent replicates, and n = number of worms taken for behavioral study.

We ablated PVD neurons on one or both sides (PVDL and PVDR, Fig 1E-F) at the L4 stage to assess the role of these neurons in harsh touch response. The percentage of worms responding to harsh touch was significantly reduced when both the PVD neurons (PVDL and PVDR) were ablated as compared to the control (mock injury) (Fig. 1F). However, in cases of ablation on one side, only the harsh touch given to ablated side resulted in a significant reduction in response (Fig. 1F). Similarly, we found that following axotomy near the cell body (red cross, Fig. 1E,G), the percentage of worms responding to harsh touch decreased when both the PVD neurons were axotomized and on the axotomized side when one axon was cut (Fig. 1G). Surprisingly, when we dendrotomized both the major and minor primary dendrites in both sides (PVDL and PVDR, Fig. 1E,H), the harsh touch response remained unaffected (Fig. 1H). This indicated that dendrotomy does not affect the harsh touch response behavior associated with PVD neurons.

We also developed a more quantitative assessment of harsh touch response by analyzing video recordings of the worm’s response to a harsh touch stimulus. We used an Image-J plugin software (Nussbaum-Krammer et al., 2015) that tracks the worm’s speed (Fig. S1A). The speed of the locomotion increased significantly following the delivery of harsh touch (Fig. S1B). The Day 1 and Day 2 adults also showed a similar increase in speed upon harsh touch (Fig. S1B). We defined the harsh touch response index (HTRI) as the ratio of the worm’s speed after to before the harsh touch delivery (Fig S2A). As expected, the HTRI value in the *mec-3* mutant is significantly lower than in the wild type (Fig S2 C). On the other hand, the HTRI was unaffected in *mec-4* mutant worms compared to wild-type worms (Fig S2 C). Additionally, HTRI levels were significantly lowered after PVD ablation or axotomy (Fig S2 D-E). This decline was specific to the injured side (Fig S2 D-E). Dendrotomy of both the PVD neurons did not affect the HTRI value (Fig S2 F).To answer the question that whether the dendritic morphology may play any role in harsh touch response or not, we checked the harsh touch response for the mutants that have a developmental defect in the formation of the dendritic arbor of PVD neuron such as *hpo-30(0)* and *tiam-1(0)*(Zou et al., 2018, Tang et al., 2019). The harsh touch response was unaffected in these mutants (Fig 1D) saying that the dendritic morphology may not be playing any significant role in harsh touch response. This indicated that harsh touch sensory modality may not be entirely dependent on the dendrites of PVD neurons.

### The dendrotomy on PVD affects proprioception behavior

As the harsh touch response behavior was not affected by dendrotomy, we speculated that the proprioception behavior might be affected by dendrotomy. The dendrites release the neurotransmitter NLP-12 to the neuro-muscular junction that modulated the sinusoidal wave pattern formed by the locomotion of the worms(Tao et al., 2019). The amplitude and wavelength of the sinusoidal waveform made by a freely moving worm on the bacterial lawn (Fig. 2A) have been described as the hallmarks of proprioceptive behavior in worms (Albeg et al., 2011, Tao et al., 2019). These parameters are often altered in mutants missing higher-order dendritic branches in PVD neurons (TAO et al. 2019, (Albeg et al., 2011). We used these parameters to test if this proprioception behavior was changed after dendrotomy. We took images of the sinusoidal traces made by the moving worms on the OP50 bacterial lawn on NGM plates (Fig. 2A). The amplitude, wavelength, and bending angle were computed from the sinusoidal trajectory (Fig 2A). The *mec-3, hpo-30,* and *tiam-1* mutants, which have reduced the number of higher-order branches in PVD (Sundararajan et al., 2019, Zou et al., 2018, Tang et al., 2019), exhibited a decrease in all postural measurements (Fig S2A-C). The mutant for *mec-10* which codes for DEG/ENaC channels also showed reduced values in these parameters (Fig S2 A-C) as shown before (Tao et al., 2019). We found that the amplitude and wavelength values at Day 1 and Day 2 adult stage were significantly higher than at larval stage L4 (Fig S2 D-F) most likely due to an increase in the length of the worm body which is 950 microns at L4 stage to 1.2 millimeters at Day 1 stage. These parameters were comparable in Day 1 and Day 2 stages. (Fig S2 D-F). Therefore, we decided to test the behavioral changes due to dendrotomy at Day 1 adult stage.

**Figure 2:**
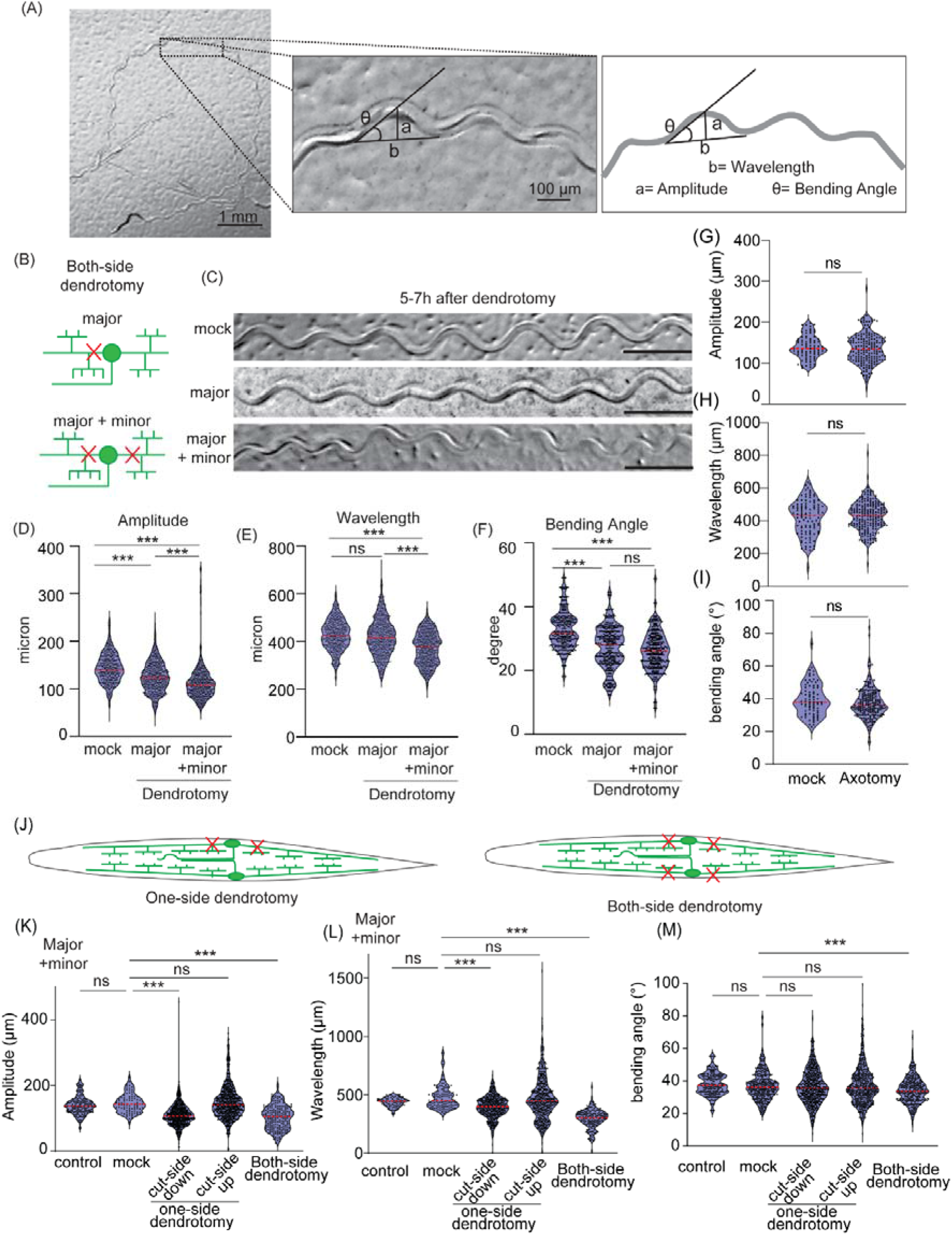
Dendrite injury in PVD results in defective posture of the worms. (A) The worm’s trajectory is shown into the OP50 bacterial lawn after a movement for 3 to 5 miunites duration. The magnified picture and schematics displaying the postural parameters calculated as amplitude labelled as ‘a’, wavelength labelled as ‘b’, and bending angle labelled as ‘θ’ measured from each wave are also shown. (B) Schematics representing the cutting experiments including the dendrotomy of ‘major dendrite only’ and dendrotomy of both ‘major and minor primary dendrites’ of both of the PVD neurons (PVDL and PVDR) of the worm is shown. (C) Trajectories at 5-7 hours after dendrotomy in mock, dendrotomy in the major dendrite (both PVDs), and dendrotomy in the major and minor dendrite (both PVDs) in *wdIS52 (pF49H12.4::GFP)* background worms are shown. The scale bar represents 500 microns. (D-F) Posture characteristics such as amplitude (D), wavelength (E), and bending angle (F) in *wdIs52 (pF49H12.4::GFP)* worms are plotted in mock and dendrotomy paradigms mentioned in B-C, 25<n<32, N=3. The absolute data were displayed, and the violin plots represented the median (red line) and population distribution. (G-I) Posture characteristics such as amplitude (G), wavelength (H), and bending angle (I) in *wdIs52 (pF49H12.4::GFP)* worms are plotted in mock and axotomy (both PVDs) conditions done at Day1 old worms, 13<n<25, N=3. The absolute data were displayed, and the violin plots represented the median (red line) and population distribution. (J) Schematics of one-side dendrotomy (PVDL or PVDR) and both-side dendrotomy injury where both major and minor dendrites are injured. (K-M) the quantification of posture parameters i.e. amplitude (K), wavelength (L) and bending angle (M) were measured and plotted for control, mock, one-side dendrotomy (cut-side down i.e. the injured PVD neuron is touching the agar media and cut-side up i.e. injured PVD neuron is not touching the agar media), and both-side dendrotomy at 8h after injury. The absolute values are plotted with red dotted line representing the median of the population data. 27<n<30, N=3. The statistical analysis for D-F,K-M is one-way ANOVA with Tukey’s multiple comparisons, G-I, student t-test, with p < 0.05*, 0.01**, and 0.001***, ns not significant. N stands for the number of independent replicates, and n stands for the number of events.

We performed two types of dendrotomy experiments: (1) dendrotomy in major primary dendrites and (2) dendrotomy in both major and minor primary dendrites for both PVD (PVDL and PVDR, Fig. 2B). The trajectories of the worms that underwent dendrotomy appeared distinctly different as compared to the mock control (Fig. 2C). The postural parameters were lowered after dendrotomy of major dendrites and with major-plus-minor dendrotomy (Fig. 2D-F). The amplitude and bending angle were significantly decreased after the dendrotomy of the major primary (Fig. 2D&F). These parameters were reduced further significantly when both major and minor dendrites were cut except wavelength (Fig 2D-F). The wavelength parameter was only perturbed with dendrotomy of major and minor dendrites where it was significantly reduced as compared to the mock injured or major dendrite dendrotomized (Fig 2E). To see whether the defect in sinusoidal posture after laser damage is a secondary consequence of locomotion defect, we evaluated the baseline speed of the dendrotomized worms (Fig. S2G). The speed of the worms remained unaltered post-dendrotomy (Fig. S2G). The gentle touch response was also unaffected in these worms (Figure S2G, lower panel). Dendritic contribution in proprioception was further confirmed when axotomized worms did not show any perturbation in the posture parameters (Fig. 2G-I). When the PVD neurons were axotomized, we found no change in the postural parameters as compared to the control (Fig. 2 G-I).

To ascertain if the dendrotomy-related drop in proprioception is specific to the transgenic strain, we repeated this experiment in another transgenic strain expressing mScarlet diffusible reporter in PVD neurons (Figure S2J-O). The amplitude and bending angle both exhibited a reduction after dendrotomy of major as well as major-plus-minor dendrite in this strain (Fig. S2M-O). Whereas, the wavelength was dramatically shortened when both major and minor dendrites are cut (Fig. S2N). We further assessed the relative contribution of PVDL and PVDR in the proprioception by injuring both major and minor dendrites on only one side (PVDL or PVDR) (Fig. 2J). The posture of the worm was correlated to the PVD neuron in contact with the substratum i.e. when the PVD with injured major and minor dendrites was in contact with the NGM plate, we observed a defect in the proprioception (Fig. 2 K-M). Both the amplitude as well as wavelength parameters were affected in the one-side dendrotomy as compared to two-side dendrotomy experiments (Fig. 2 K-M).

This indicated that axons and dendrites are differentially regulating the harsh touch response and proprioception, respectively.

### Axon regrowth of PVD neurons restores the impairment in harsh touch response

Previous studies have elucidated that axon regeneration from the injured cut stump leads to functional recovery in various model systems (Laha et al., 2017, Becker et al., 1997, Rasmussen and Sagasti, 2017, Basu et al., 2021). As PVD neurons also show axonal regeneration (Brar et al., 2022), we investigated whether the deficits in the harsh touch response due to PVD axotomy would be restored in subsequent time points when this axon regenerates. After the axotomy near the cell body (red arrow, Figure 3A), we saw a retraction of the injured tip, followed by new neurite formation and regrowth towards the ventral nerve cord (Fig. 3A). The proportion of worms responding to harsh touch at 24 hours after axotomy was higher than at 8-10 hours after axotomy in both one-side and two-side axotomy paradigms (Fig. 3B). Similarly, HTRI values also showed the recovery of HTRI at 24 hours after axotomy and is side specific recovery (Fig. 3C).

**Figure 3:**
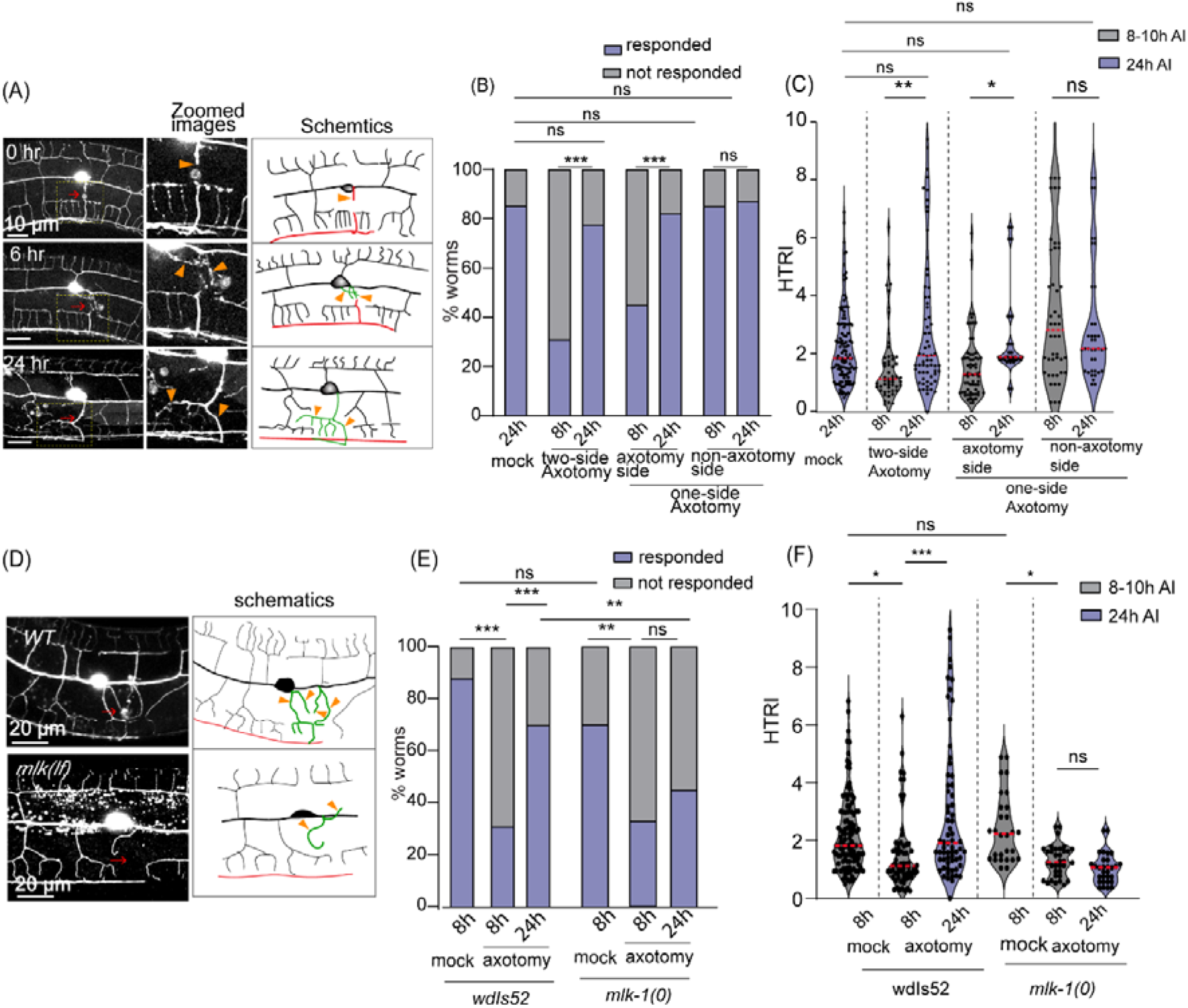
Axon regeneration leads to recovery in Harsh touch defect. (A) Confocal images along with the schematics of of PVD neuron is shown at 0 hour, 6 hour and 24 hour after axotomy representing the axon in red, new protrusions in green traces, and orange arrowheads indicate the new regrowing branches. The red arrow mark the injury site. (B-C) Quantification of harsh touch as percentage of worms responding (B) and Harsh touch response indices (HTRI) (C) of mock (24h), axotomized-side and non-axotomy-side in one-side axotomy (8h and 24h) and two-side axotomy (8h and 24h) in *wdIs52 (pF49H12.4::GFP)* worms. 24<n<32, N=3 (B). 21<n<35, N=3 (C). (D) The confocal images of wild-type and *mlk-1(0)* in *wdIs52* background are shown along with the schematics representing the axon in red, regrowing branches in green, and orange arrowheads represent the regrowing branches at 24h after injury. The red arrow mark the site of injury. (E-F) Worms responding to harsh touch as percentage (E) and Harsh Touch Response indices (F) were quantified and plotted for mock at 8h, axotomy at 8h, and axotomy at 24 h after axotomy of both the PVD neurons in *wdIs52* and *mlk-1(0); wdIs52* background. 18<n<35, N=3 (E). 18<n<35, N=3 (F). The statistical analysis B, and E, is Fisher’s exact two-tailed test, for C, and F, is one way ANOVA with Tukey’s multiple comparison test with p<0.05*, 0.01**, and 0.001***. Violin plots represent the median and population distribution. ns stands for not significant, N stands for the number of independent replicates, and n stands for the number of worms taken for analysis.

DLK/MLK pathway is a critical regulator of axon regrowth following injury (Yan et al., 2009, Nix et al., 2011, Brar et al., 2022). To study if DLK/MLK mediated axonal regrowth is required for functional recovery, we assessed the harsh touch response in the *mlk-1* mutant (Fig. 3D). After axotomy in the wild-type background, we noticed regrowth towards the ventral nerve cord (Fig. 3D) as seen before (Brar et al., 2022).

The proportion of the worms responding to the harsh touch also showed an improvement at 24h. The regrowth was diminished in the *mlk-1* mutant (Figure 3D) and we noticed that the percentage of *mlk-1(0)* worms responding to harsh touch at 24 h did not improve as compared to 8 h after injury (Figure 3E-F). The HTRI values also showed a recovery from 8 h to 24 h after injury in the wildtype worms while this recovery was absent in the *mlk-1(0)* (Fig. 3F). This indicated a strong correlation between the axon regeneration to the recovery of harsh touch sensation in the PVD neurons. This observation got further strengthened by the measurement of harsh touch response in the wild-type worms with the ablated PVD neurons (Fig. S3 A-B). In these worms, the harsh touch response did not improve at 24 hours post-ablation as observed by the percentage of worms responding to harsh touch and HTRI values as compared to their respective 8 h readouts (Fig. S3 A-B). This was consistent irrespective of the injured side with respect to the harsh touch stimulus (Fig. S3 A-B). These results suggest that the axon regeneration in PVD neurons enables functional recovery of harsh touch sensation in these neurons and this recovery is blocked when axon regeneration is perturbed.

### Functional restoration during dendrite regeneration correlates with successful events of arborization and fusion

Our previous study showed that the PVD neurons regenerated their dendrites following a laser-assisted injury (Brar et al., 2022). However, it is unclear how the proprioception or the sinusoidal movement of the worm changes with regenerating dendrites. Dendrotomy of either one or both PVD neurons resulted in a drop in the amplitude and wavelength of the sinusoidal traces made by the moving worm (Fig 4A-B). Qualitative inspection of the traces of dendrotomized worms showed a marked improvement at 24 h post dendrotomy with lesser irregularities in the sinusoidal movement as compared to the traces observed at 8 h post dendrotomy (Fig. 4C-E). All the parameters of proprioceptive behavior including amplitude, wavelength, and bending angle were significantly increased at 24-hour post-dendrotomy as compared to their 8 h counterparts (Fig 4C-E).

**Figure 4:**
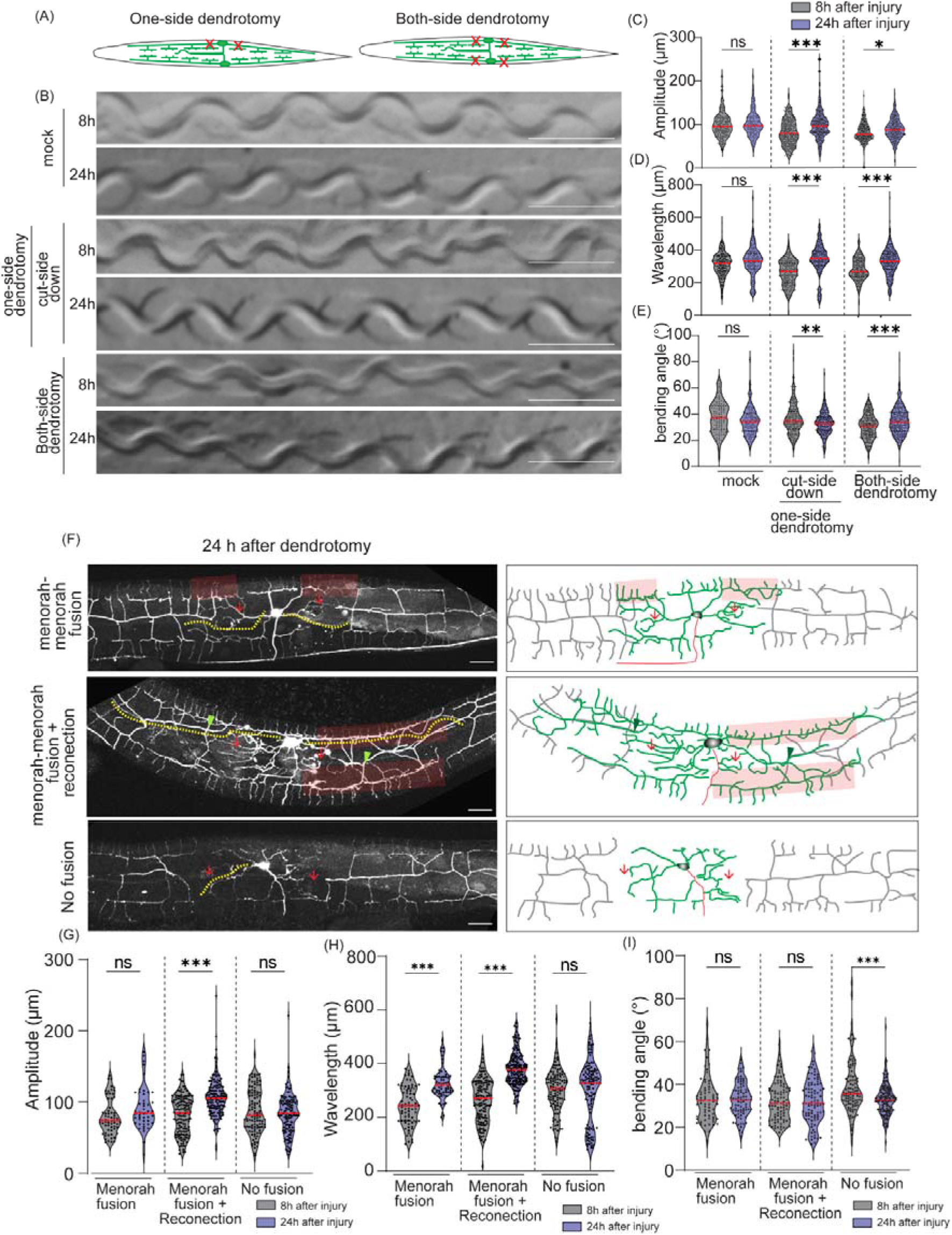
Dendrite regeneration in PVD neurons leads to recovery in posture defect of worms. (A) Schematics representing the one-side and both-side dendrotomy where both the major and minor dendrites were injured. (B) The images of the worm trajectories on bacterial lawn are shown at 8h and 24h after injury in mock, one-sided dendrotomy (cut-side down where the injured PVD is touching the agar media) and both-side dendrotomy conditions. The scale bar represents 500 microns. (C-E) Quantification of amplitude (C), wavelength (D) and bending angle (E) at 8h and 24h after injury in mock, one-side dendrotomy (cut-side down i.e., injured PVD neuron is touching the agar media) and two-side dendrotomy are plotted with the absolute values in a violin plots. The red dotted lines represent the median of population occurrence. 27<n<30, N=3. (F) Representative confocal images along with the schematics at 24 hours after dendrotomy with the menorah fusion highlighted by red faded box, the green arrowheads marking the reconnection phenomena and the red arrows represent the site of injury. The regenerated dendrites are represented in green, the distal part in grey and axon is represented in red in the schematic. The yellow dotted lines represent the longest regenerating dendrite corresponding to the territory covered by regenerated dendrites. Three different cases are shown i.e., menorah fusion, menorah-menorah fusion plus reconnection and no fusion events. (G-I) The quantification of postural parameters i.e., amplitude (G), wavelength (H) and bending angle (I) are shown at 8hours and 24 hours after dendrotomy which are classified in three different cases which are menorah fusion, menorah fusion plus reconnection and no fusion are plotted. 15<n<22, N=3. The statistical analysis for C-E, G-I is Tukey’s multiple comparison test, with p<0.05*, 0.01**, and 0.001***. The violin plots represent the median and population distribution. ns stands for not significant, N stands for the number of independent replicates, and n stands for the number of worms taken for analysis.

Dendrite regeneration in the PVD neurons is accompanied by regrowth from the severed end, reconnection between distal and proximal counterparts, and fusion of the higher-order branches (Brar et al., 2022). The higher-order branches often fuse with each other to bypass the disjointed primary dendrites termed menorah-menorah fusion (Oren-Suissa et al., 2017, Kravtsov et al., 2017, Brar et al., 2022). In these experiments also we noticed similar branching with reconnection (green arrowheads) and menorah-menorah-fusion (red faded box) events (Fig. 4F). The regrowing dendrites span the gap created by the injury. This territorial expansion is estimated by the length of the longest neurite measured from the cell body (Yellow dotted line, Fig. 4F).

Due to the pleiotropic nature of the regenerative growth of the dendrites, it is difficult to ascertain if the pattern of regrowth attributes some advantage in the recovery of proprioception. We divided the events into three categories, “menorah-menorah fusion”, “menorah-menorah fusion plus reconnection”, and “no fusion” (Fig. 4F). Based on the amplitude, wavelength, and bending angle measurements of the sinusoidal movement, recovery was observed in cases with menorah fusion (Fig. 4G-I). In cases with menorah fusion plus reconnection, both amplitude and wavelength values were significantly improved at 24h as compared to 8h post-dendrotomy (Fig. 4G-H). On the other hand, we did not observe any functional recovery in cases of “no fusion” events (Figs. G-H). Interestingly, the bending angle did not change during dendrite regeneration except in cases of no fusion and reconnection which showed a decrease (Fig I). This could be due to a decline in neuronal and muscle vigor in the absence of a robust or interconnected dendritic arbor. This is in line with functional recovery correlated to axon fusion events during regeneration in PLM gentle touch neurons (Basu et al., 2017, Abay et al., 2017).

The proprioception parameters were also checked for both side ‘major’ and ‘major plus minor’ dendrotomy experiments in worms with different transgene labeling PVD neurons with *mscarlet* fluorophore. The trajectories on the OP50 lawn showed a visible recovery 24 hours after injury (Figure S4 D). The regeneration events including reconnection and menorah fusion were also seen in these worms 24 hours after injury (Figure S4 E). We found out that the affected parameters i.e., amplitude and bending angle in case of major dendrotomy and all three parameters in case of major plus minor dendrotomy experiments showed a recovery at 24 hours after injury (Figure S4 F-H).

When these proprioception parameters were correlated with the territory length of regenerated dendrites, we found out that both the amplitude and wavelength were positively correlated with the territory length except the bending angle (Fig S4 A-C). These indicated that based on recovery of dendritic territory, the worms could regain their posture which is also facilitated by the fusion in higher order dendrites.

## Discussion

For faithful transduction of information in functional neuronal circuits, both dendrites and axons within an individual neuron play important roles (Blockus and Polleux, 2021). During the massive injury in the spinal cord, both axons and dendrites can get injured. In case of ischemia, stroke, and epilepsy, the dendritic arbors are prominently affected (Risher et al., 2010, Gao et al., 2011, Swann et al., 2000). The nerve graft-mediated stimulation of neurite regeneration in the injured spinal cord (Kumamaru et al., 2019) would also involve re-specification of the dendrites from the stem cell in the graft. Therefore, functional rewiring after neuronal injury would require correct regrowth and integration of the regenerated dendrites to its presynaptic neuron or sensory organ. Although recent studies have shown that targeted injury on dendrites induces regrowth from the injured tip (Thompson-Peer et al., 2016, Stone et al., 2014, Brar et al., 2022), its functional consequence in adulthood is not clear. In this study, we systematically quantified the functional decline of both touch response and proprioception following axon and dendrite injury on PVD neurons of *C. elegans*. Interestingly, we found that injury to axon and dendrite affects touch response and proprioception, respectively. This is consistent with the recent finding that dendrites and axons exclusively regulate two sensory modalities (Tao et al., 2019).

Moreover, we showed that axon regeneration leads to functional recovery in touch response in a DLK-1/MLK-1 pathway-dependent manner. On the other hand, dendrite regeneration leads to recovery in body posture defects caused due to dendrotomy. The dendrite regeneration is independent of axon regeneration pathways such as DLK-1/MLK-1 (Stone et al., 2014, Brar et al., 2022), and it requires a unique mechanism involving GTPase and GEF activity (Brar et al., 2022). Dendrite regeneration in *Drosophila* da9 neurons is correlated to the functional recovery of nociceptive function in the larval stage (Hertzler et al., 2023). Moreover, recent studies in fish indicated a regeneration response after the injury to the dendrites of spinal motor neurons (Stone et al., 2022). In the mouse brachial plexus injury model, it is seen that dendrite regeneration is controlled by Rho GTPase (Li et al., 2023). Therefore, our findings in nematode highlight the functional importance of the dendrite regeneration process and its underlying molecular mechanisms.

We see that the fusion process between the proximal and distal dendrites highly correlates to the functional recovery in the body posture during locomotion. Similar findings were seen in case of repair of injured axons in earthworm, cray fish, Leech, Aplysia and nematode (Birse and Bittner, 1976, Hoy et al., 1967, Macagno et al., 1985, Bedi and Glanzman, 2001, Ghosh-Roy et al., 2010, Neumann et al., 2011). In *C elegans*, the fusogen molecules such as EFF-1 and AFF-1 mediate fusion process in axons and dendrites (Ghosh-Roy et al., 2010, Neumann et al., 2015, Oren-Suissa et al., 2010, Kravtsov et al., 2017). Here, we found that the recovery in some of the body posture parameters also correlates to the length of the territory covered by the regenerated dendrites.

Overall, our study using PVD model establishes that dendrite regeneration can restore the lost sensory function due to injury in adulthood (Fig. 5). It also establishes that axon and dendrite regeneration is both molecularly and functionally distinct (Fig. 5). It further underscores the importance for studying a detailed molecular mechanism controlling the dendrite repair process.

**Figure 5.**
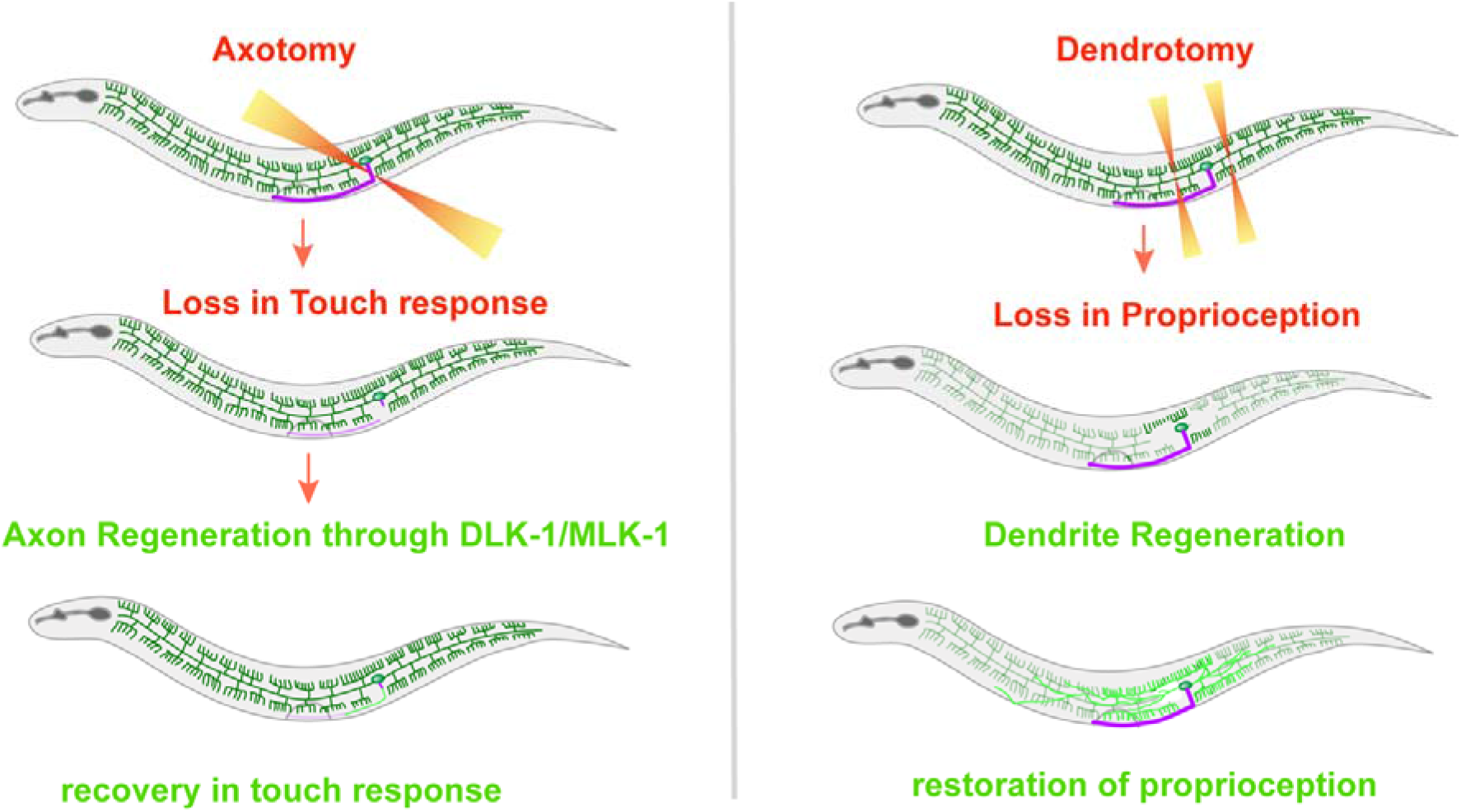
Differential regulation of sensory modalities of PVD neuron by its axon and dendrites. The model depicts the contribution of the axonal and dendritic compartments in regulation of the nociceptive harsh touch and proprioception in worms, recpectively. Loss of axon and dendrite cause a decline in harsh touch and posture, respectively which is regained when these compartments regenerate. Conditions regulating the axon and dendrite regeneration in PVD neurons also correlate well in anatomical and functional aspects of this neuron.

## Materials and Methods

### Strains of *C. elegans*

*C. elegans* strains were grown and maintained at 20°C on the nematode growth medium (NGM) plates seeded with the OP50 bacteria (Brenner, 1974). The loss of function mutants are represented as (0), for example, the *dlk-1* loss-of-function allele *tm4024* is mentioned as *dlk-1(0)*. Unless otherwise specified, the mutants used in this study are deletion or substitution mutants (Table S1). The Caenorhabditis Genetics Centre (CGC) provided these strains, which were genotyped using the appropriate genotyping primers.

### Dendrotomy and axotomy

Dendrotomy and axotomy experiments were performed on *C. elegans* using a Bruker® system equipped with a SpectraPhysics® femtosecond multiphoton laser system and an Olympus® 60X/0.9NA water objective. The neurite imaging and severing were performed using 920nm and 720nm wavelengths, respectively. Worms were mounted with cover glass on 5% agarose pads with a drop of polystyrene beads (Polysciences 00876-15) as a friction raising agent. In some experiments Levamisole hydrochloride (10mM) (Sigma L0380000) was used as a paralyzing pharmacological agent (cat No. 2855-25).

Usually for dendrotomy, two laser shots were delivered to create a big gap in the primary dendrites (Figure 1E) of PVD as described before (Brar et al., 2022). The first laser-shot was delivered at the junction of the first secondary (9-10 μm from the cell body), and the second at a 10- μm distance from the first cut site. The axon of PVD neuron was severed at 5 μm away from the cell body, resulting in a noticeable gap (Figure 1E). As control, mock injuries were carried out away from the PVD neurites with equivalent number of laser shots to other cohorts of injuries performed in any particular experiment. The injured worms were then recovered to OP50-seeded NGM plates using a mouth pipette. To assess the behvior of the injured PVD, injury paradigms of ablation and axotomy involved either one PVD (PVDL or PVDR) or both (PVDL and PVDR) are mentioned in the respective figure-panels. For dendrotomy, one paradigm involved injuring major and minor dendrites of either of one PVD neuron (PVDL or PVDR) or both the PVDs (Fig 2J). Second dendrotomy paradigm involved injuring either major dendrite or major and minor dendrites of both the PVD neurons to understand role of major dendrite specifically (Fig 2B).

### Imaging

To measure the degree of regeneration, injured worms were imaged at 24 hours (h) after the injury. The worms were immobilized and plated on 5% agarose (Sigma®) pads in 10mM levamisole hydrochloride (Sigma®) media. The worms were scanned using a Nikon® A1R confocal system and a tile imaging module with a 60X/1.4NA oil objective to measure the degree of regeneration.

### Dendrite and axon regeneration analysis and quantification

The proximal and distal ends of injured PVD dendrites can fuse after injury (Kravtsov et al., 2017). The overlaps between the regenerating proximal primary dendrite with distal primary dendrite (Fig 4F, green arrowhead) or distal menorah (Fig 4 F, green arrowhead) were classified as reconnection phenomena as also described before (Brar et al., 2022).. The menorahs having more than one secondary dendrite (Fig 4F, faint red rectangular boxes) were considered as menorah-menorah fusion and the percentage of PVD neurons showing menorah-menorah fusion was calculated and plotted.

Analysis of the axon regeneration was carried out by visual observation and quantitative information was obtained using various image analysis modules of ImageJ®. Neurites were traced and their lengths were quantified using the Simple Neurite Tracer plugin of ImageJ® (Schindelin et al., 2012). Neurite development from the axon’s split point was classed as regenerative growth including the ectopic branches that originate from the cell body, the nearby dendrite, and the overall branch length. To collect statistical data, ImageJ® quantifications were further examined in Excel and Graphpad®.

### Behavioral Analysis of Harsh touch response

In this assay, we used a 0.15 mm thick platinum wire in a cuboid form, which has previously been used to provide a harsh touch of 100-200 N force (LI et al. 2011). The worms were allowed to roam freely on the NGM medium plates and harsh touch was provided near the vulva. The assay involved providing the harsh touch by the platinum wire on both left and right sides of the worm, categorized separately for either side and later correlated for the injured (ablated/axotomized/cut side down) and uninjured (unablated/non-axotomy/cut side up) side in cases of one PVD injured (PVDL or PVDR).

The worms displayed an escape response with increased velocity or changed direction or both. The live-imaging of the touch response was done using a Leica stereoscope (LEICA M165 FCM165FC) with a camera (LEICA MC120 HD) attached. The recorded videos were analyzed using the wormtracker plugin in the ImageJ® Fiji program (Nussbaum-Krammer et al., 2015). The proportion of worms displaying an escape response after the harsh touch to the total number of worms assessed was quantified as the percentage of worms “responded”. Similarly, the worms without any escape response following harsh touch have been classified as percentage of worms “not responded”.

To quantify the response to harsh touch, each of the worms was recorded for 30 seconds before and after the harsh touch. The quantitative strategy was carried out by utilizing the ImageJ® plugin ‘wrmtrck’(Nussbaum-Krammer et al., 2015). For each worm, before and after the harsh touch videos were segregated and analyzed. After transforming the videos into 8-bit videos and thresholded, the wrmtrck plugin was used with parameters such as (minsize=1000, maxsize=19999, maxvelocity=800, maxareachange=900, mintracklength=2, bendthreshold=1, binsize=0, showpathlengths, showlabels, showpositions, smoothingrawdata=2, benddetect=0, fps=10, backsub=0, threshmode=Otsu, fontsize=16). The average speed values were calculated from the videos of 30 second duration before and after harsh touch. This ratio of speeds after the harsh touch to before the harsh touch was termed a harsh touch response index or HTRI (Fig. S1A).

### Behavioral Analysis of the Posture of Worm

The worms were put onto the 3 days old seeded OP50 lawn using an eyelash pick and were let to freely crawl for 5 mins. The proprioception assay involved placing the worm with their left and right sides on the OP50 lawn, categorized separately for either side and later correlated for the injured and uninjured side in cases of one PVD injured (PVDL or PVDR). For the posture assay, still images of the sinusoidal trajectories on OP50 lawn were acquired with a Leica camera (LEICA MC120 HD) after the worms were alllowd to move for 5 mins The posture parametersincluded analysis of the amplitude, wavelength, and bending angle of each sinusoidal wave manually determined using Fiji® ImageJ software (Fig 2 A). The amplitude (a) was measured using a line segment measuring the height of the peak (Fig 2A, a). The wavelength (b) was measured using a line segment that connects the two minima (Fig 2A, b). The bending angle (θ) was measured using the angle tool of Image J software measuring the angle of the trajectory from the normal (Fig 2A, θ). The dataset to be utilized for the statistical analysis includes pooled values of all worms in a certain group. To straighten the traces for representation, the Straighten tool of ImageJ is used based on a segmented line with line width 50 spanning the entire width of the trajectory (Fig 2 A).

### Double-blinded analysis of worm behavior

We double blinded the behavioral assays in this study by involving two researchers for a behaviour experiment after dendrotomy or axotomy. The worms were injured using 2-photon lasers by one researcher and pseudolabeled. The other researcher then carried out the behavioral assay and analysis on these pseudolabeled worms. For example, dendrotomy experiments were done in three groups, mock, dendrotomy in major, and both minor-plus-major of one side of PVD neuron by one researcher and each cohort was pseudolabeled. Later behavioral studies and regeneration imaging were performed on these worms by another researcher, unaware of the conditions of these cohorts. Each of these cohorts were then assessed as per the categorization of pseudolabels. Behavior such as proprioception and harsh touch was done on these worms at two time points i.e. 8 hours and 24 hours. The behavioral observations were made for both left and right sides of the worm and categorized separately. Later on, the regeneration pattern at 24 hours was imaged for the same worms using confocal microscope. The information on behavioral and regeneration patterns were saved for each worm and then correlated for further studies. Based on the side of the worm receiving the harsh touch or placed on the substratum, behavioral readouts were classified for the injured (ablated/axotomized/cut side down) and uninjured PVD (unablated/non-axotomy/cut side up) in cases of only one PVD injured (PVDL or PVDR).

### Statistical analysis

To prepare statistical analyses, the GraphPad Prism® application (Prism 8 V8.2.1) was utilized. Two samples were evaluated using the unpaired two-tailed T test. To do statistical analysis on a large number of samples, one-way ANOVA with Tukey’s multiple comparisons test was carried out. Two-tailed Chi-square to compare population data, Fisher’s exact contingency test was employed to compare fraction values for each sample. For each plot, the legends show the number of samples (n) and biological replicates (N). Significance values of p < 0.05*, 0.01**, and 0.001*** obtained through statistical analysis are mentioned in the graphs.

## Supporting information

Supporting figures with captions

## Acknowledgments

We thank National BioResource Project (NBRP), Japan, and Caenorhabditis Genetics Center (CGC) for strains. We thank Smriti Bhardwaj for making the P*ser2prom4[4.1kb]::mScarlet* [*shrEx472*] transgenic worm. This work was supported by the National Brain Research Centre core fund from the Department of Biotechnology, The Wellcome Trust DBT India Alliance (IA/I/13/1/500874), and a grant from the Science and Engineering Research Board (SERB: CRG/2019/002194) to A.G-R. The Caenorhabditis Genetics Centre is supported by the National Institutes of Health Office of Research Infrastructure Programs (P40 OD010440).

## Author Contributions

H.K.B, S.D, D.P. and A.G-R designed experiments. H.K.B, and D.P. performed experiments and analyzed data. S.D. and P.S. perfomed double-blinded experiments in behavioural assays. P.S. edited the manuscript. H.K.B, S.D and A.G-R wrote the manuscript.

