## Supporting figures with captions for "Functional recovery associated with dendrite regeneration in PVD neuron of *C. elegans*"

**List of supplementary figures**

**Figure S1, related to Figure 1:** Harsh touch response index measured in different injury conditions of the PVD neurons

**Figure S2, related to Figure 2:** Posture function of PVD neurons is correlated to its dendritic structure.

**Figure S3, related to figure 3,** Complete loss of harsh touch function due to ablation of PVD neurons.

**Figure S4, related to Figure 4:** Characteristic pattern and extent of dendrite regeneration is correlated with recovery in postural parameters following injury

**List of Tables**

**Table S1**List of *C. elegans* strains used in this paper.

**Table S2**List of strains carrying extrachromosomal transgenes used in this paper.

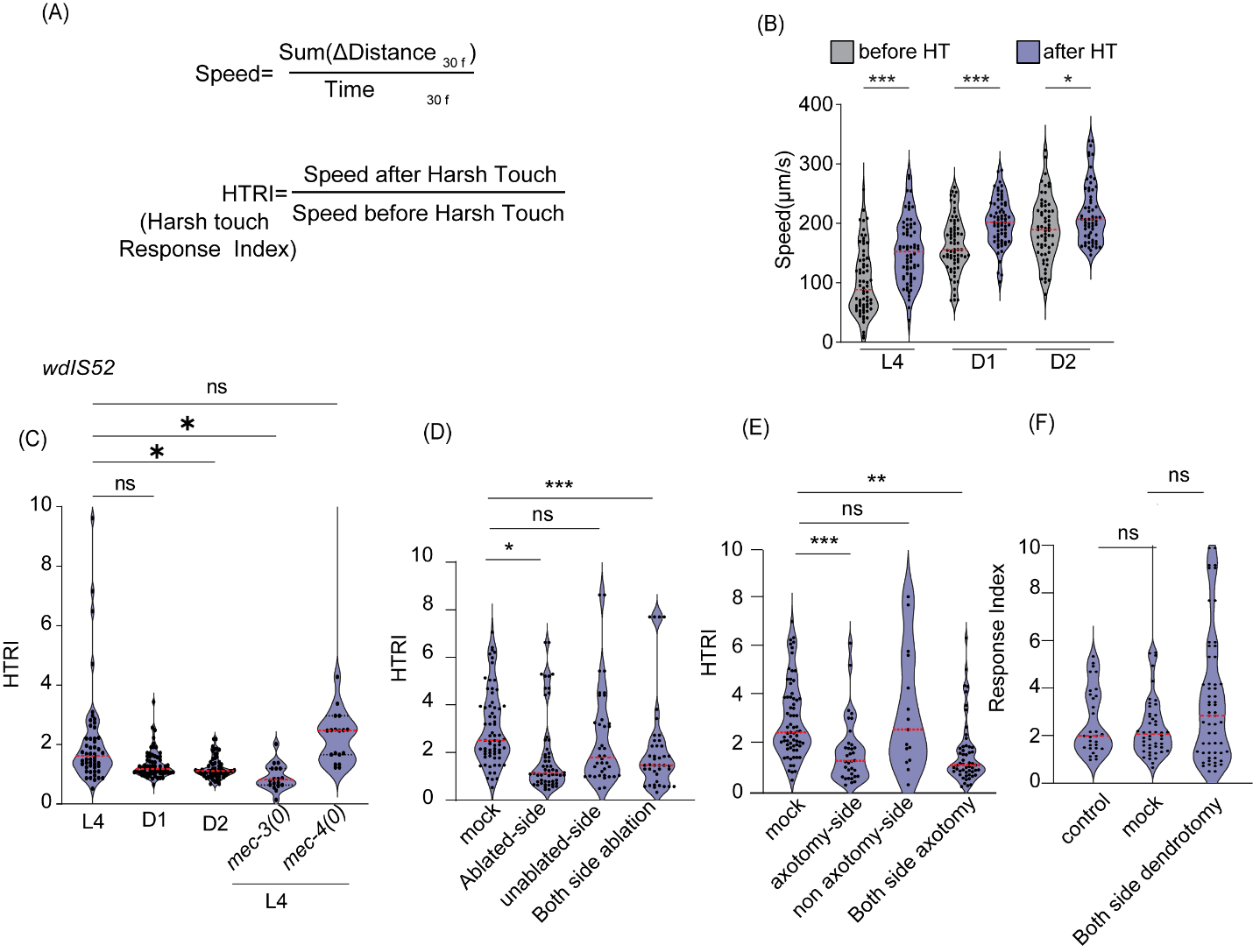

**Figure S1, related to Figure 1: Harsh touch response index measured in different injury conditions of the PVD neurons**

(A) The formula of speed and Harsh touch response index (HTRI) is shown which was used in the analysis of injury experiments. The speed of the worm was measured considering 30 frames before and after harsh touch which was labelled as speed before harsh touch and speed after harsh touch, respectively. The ratio of these speeds was taken as HTRI. (B) The values of speed of uninjured worms are plotted before and after harsh touch at L4, Day 1, and Day 2 old stage worms on the NGM plates are shown in microns per second. 12<n<30, N=3. (C) Harsh touch response indices are plotted for L4, Day1, Day2, L4 (*mec-3(0)*) and L4 (*mec-4(0)*) worms in *wdIs52 (pF49H12.4::GFP)* background. The violin plots represent the median (red line) and population distribution. 12<n<25, N=3. (D-F) Harsh touch response indices are plotted for ablation (D), axotomy (E), and dendrotomy (F) experiment. Each violin plot in (D) represents mock, one side ablation (ablated-side, and unablated side) and both side ablation in *wdIs52 (pF49H12.4::GFP)* background. 15<n<25, N=3. (E) plot represents mock, one side axotomy (axotomized-side, and non-axotomy side) and both side axotomy in *wdIs52 (pF49H12.4::GFP)* background. 13<n<35, N=3 and (F) represents control (uncut), mock, and both side dendrotomy in *wdIs52 (pF49H12.4::GFP)* background. 12<n<28, N=3. The violin plots represent median (red line) and population distribution. The statistical analysis for (B-F), is one-way ANOVA with Tukey’s multiple comparisons with p value as p < 0.05*, 0.01**, and 0.001***. ns stands for not significant, N stands for the number of independent replicates, and n stands for the number of worms taken for behavioral study.

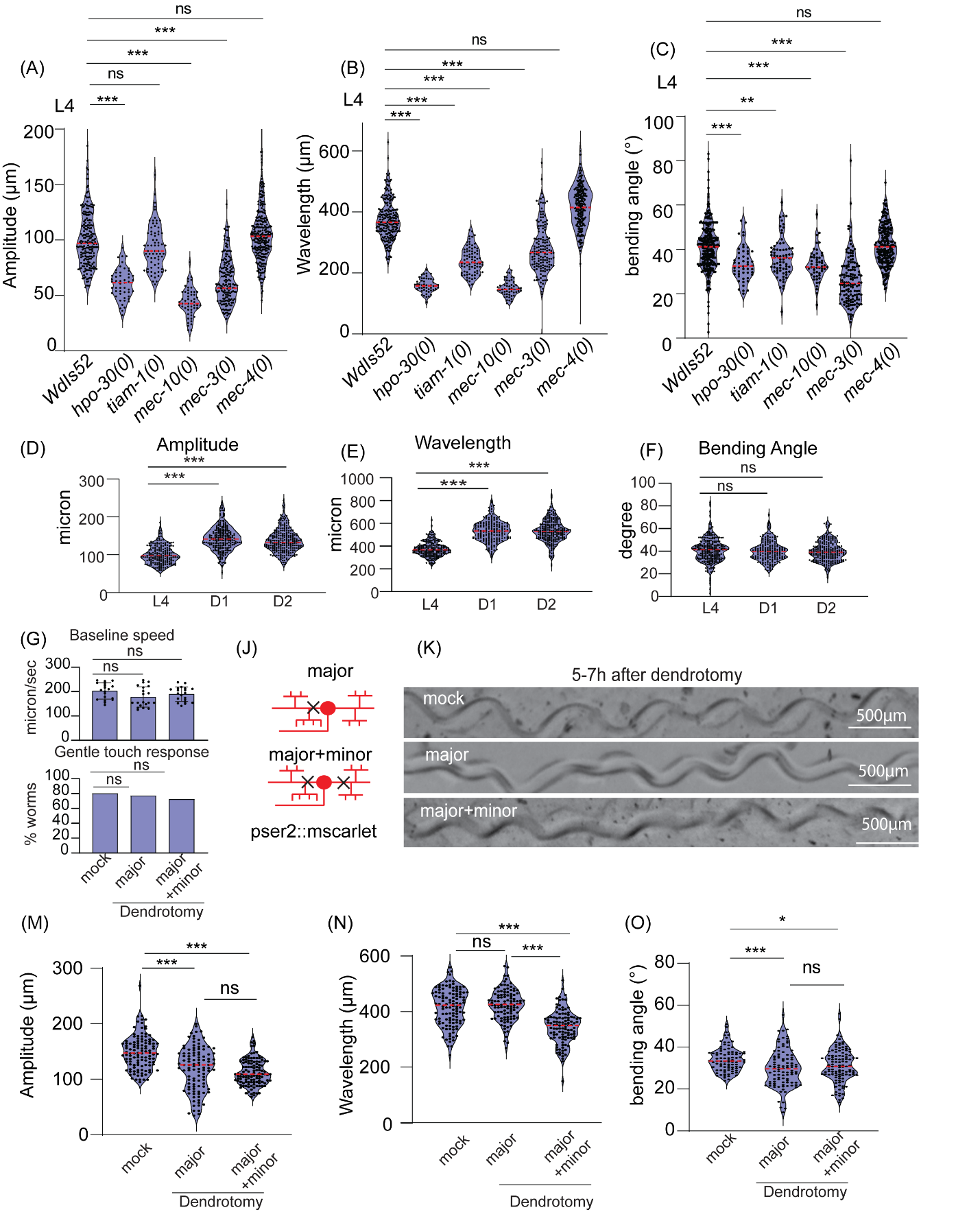

**Figure S2, related to Figure 2: Posture function of PVD neurons is correlated to its dendritic structure.**

(A-C) Posture parameters of uninjured worms i.e. amplitude (A), wavelength (B), and bending angle (C) are plotted in *wdIs52, hpo-30(0);wdIs52, mec-3(0);wdIs52, mec-4(0);wdIs52, tiam-1(0) ;wdIs52 , and mec-10(0)* at L4 stage are plotted, 25<n<30, N=3. The absolute values were plotted and the violin plots represent median and population distribution. (D-F) Posture parameters such as amplitude (D), wavelength (E), and bending angle (F) are plotted in L4, Day1, and Day2 worms respectively. The absolute values were plotted and violin plots represent median (red line) and population distribution. 15<n<25, N>3. (G) The baseline speed as well as gentle touch response were measured for the worms that undergone dendrotomy of major dendrite (both PVDs) and dendrotomy of major and minor dendrites (both PVDs) of *pser2prom3::mscarlet* worms, 13<n<25, N>3. (J) The schematics showing the type of injury performed on *pserprom3::mscarlet* worms using a 2-Photon laser. The PVD neuron is labelled in red and black cross represent the site of injury. (K) The images of trajectories at 5-7h after dendrotomy at Day 1 stage *pser2prom3::mscarlet* worms in mock, Dendrotomy in the major dendrite (both PVDs) and dendrotomy in the major and minor dendrite (both PVDs) are shown. (M-O) Posture parameters i.e. amplitude (M), wavelength (N), and bending angle (O) are plotted in mock, dendrotomy in the major dendrite (both PVDs) and dendrotomy in the major and minor dendrite (both PVDs) in Day1 *pser2prom3::mscarlet* worms. 15<n<20, N=2. The absolute values were plotted and the violin plots represent median and population distribution. The statistical analysis for A-G, M-O is one-way ANOVA with Tukey’s multiple comparisons and for (G-bottom panel) is –Fisher's two-tailed exact test with p < 0.05*, 0.01**, and 0.001***. The violin plots represent the population distribution. ns stands for not significant, N stands for the number of independent replicates, and n stands for the number of worms.

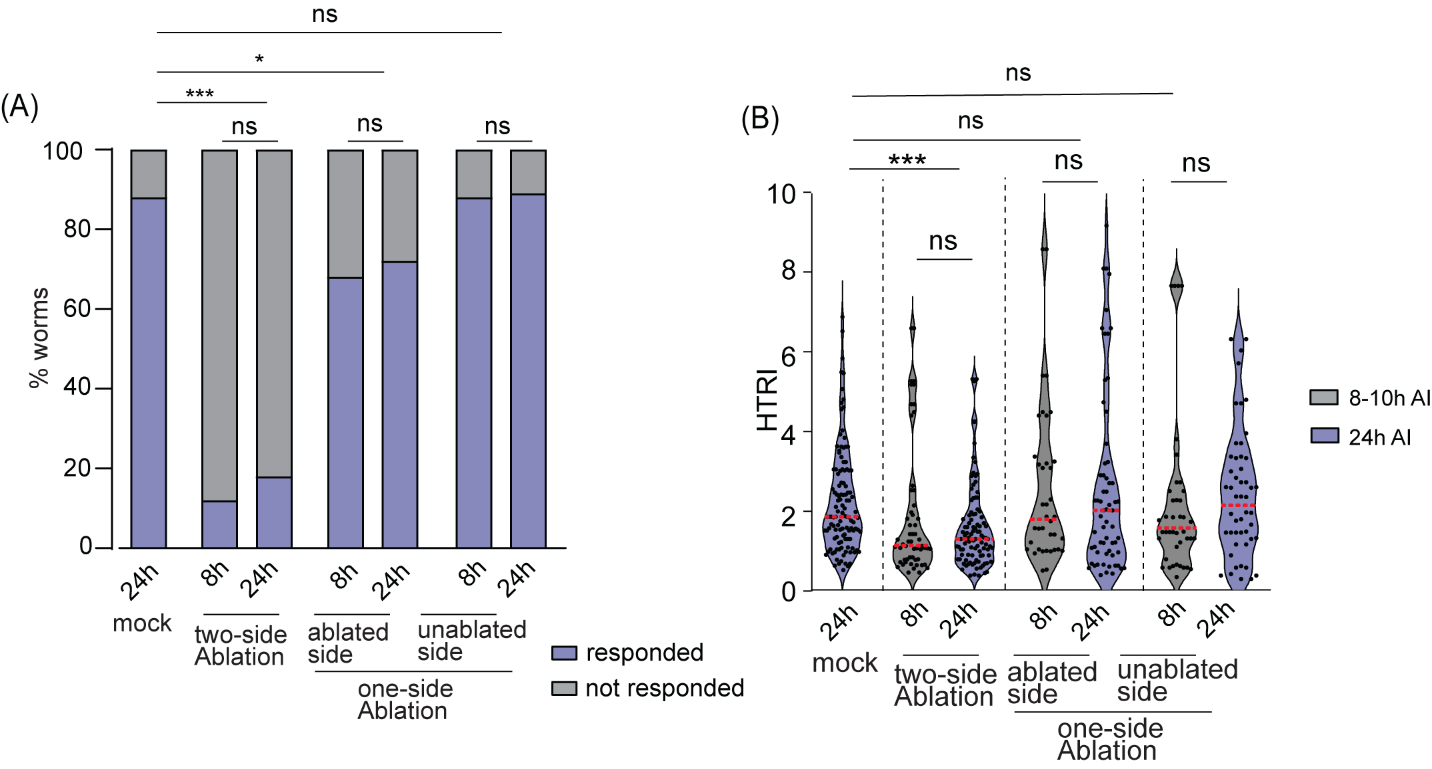

**Figure S3, related to figure 3, Complete loss of harsh touch function due to ablation of PVD neurons.**

Worms with one or both PVD ablated quantified as percentage responding to harsh touch (A) and harsh touch response indices (B) in conditions of harsh touch to mock (24h), ablated-side and unablated-side in one-side ablation (8h and 24h) and, two-side ablation (8h and 24h) after injury. 24<n<32, N=3 (A). 21<n<35, N=3 (B). The statistical analysis for A, is Fisher’s exact two-tailed test , and for B, is one way ANOVA with Tukey’s multiple comparison test with p<0.05*, 0.01**, and 0.001***. Violin plots represent the median (red line) and population distribution. ns stands for not significant, N stands for the number of independent replicates, and n stands for the number of worms taken for analysis.

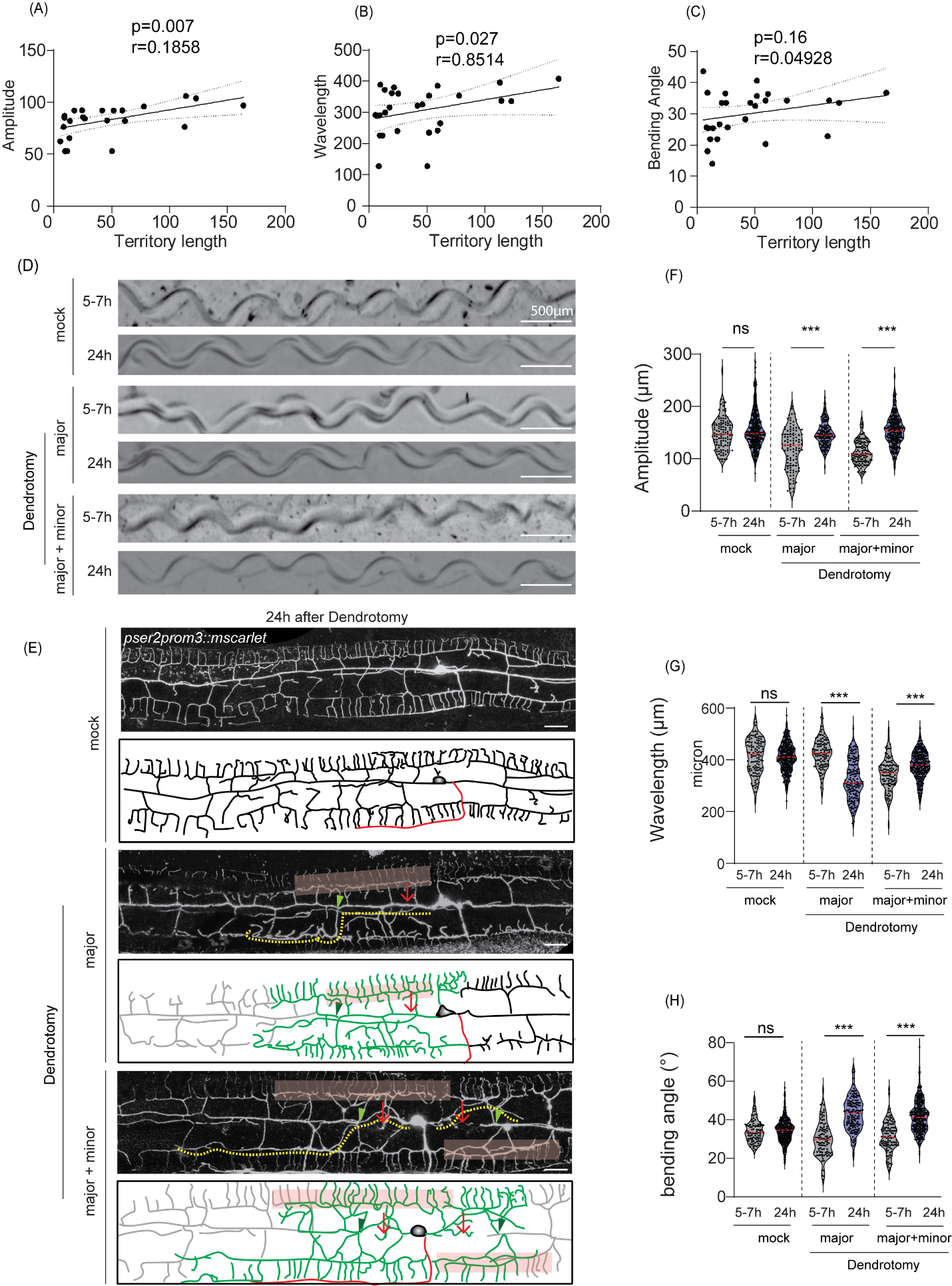

**Figure S4, related to Figure 4: Characteristic pattern and extent of dendrite regeneration is correlated with recovery in postural parameters following injury**

(A-C) The linear regression plot between various postural parameters such as amplitude (A), wavelength (B), and bending angle (C) to the territory length of regenerated dendrites. Regression analysis parameters are depicted in the plots with line of best fit, curved lines spanning the 95% confidence band and p values non zero regression fit. The cut-side down parameters were correlated with the territory length of one-side dendrotomy experiments. 10<n<32, N=2. (D) The images of trajectories at 5-7hours and 24hours after dendrotomy at Day 1 *pser2prom3::mscarlet* worms in mock, dendrotomy in the major dendrite (both PVDs), and dendrotomy in the major plus minor dendrite (both PVDs) is shown. The scale bar represents 500 microns. (E) the confocal images along with schematics representing dendrite regeneration in green, the axon in red, and the distal part in grey color. The red arrow marks the site of injury, the faint red box represents menorah-menorah fusion and green arrowheads represent primary branch reconnection. The yellow dotted line represents the territory covered. The scale bar represents 10 microns. (F-H) Posture parameters i.e. amplitude(F), wavelength (G), and bending angle (H) are plotted in mock, dendrotomy in the major dendrite (both PVDs) and dendrotomy in the major plus minor dendrite (both PVDs) in Day 1 *pser2prom3::mscarlet* worms is shown , 15<n<20, N=2. The absolute values were plotted with the median (red line) and population distribution. The statistical analysis for (A-C) is simple linear regression test with goodness of fit is calculated and its non-zero significance, (F-H) was one way ANOVA with Tukey’s multiple comparison test with p< 0.05*, 0.01**, and 0.001***. The violin plots represent the median and population distribution. ns stands for not significant, N stands for the number of independent replicates, and n stands for the number of worms taken for analysis.

| **Table S1 List of C. elegans strains used in this paper** | | |
| --- | --- | --- |
| ***C. elegans* strains** | **Source** | **Identifier** |
| *WdIs52(F49H12.4::GFP, unc119[+]) X* | CGC | NC1687 |
| *pser2prom3::mscarlet; ttx3::RFP(ShrEx472)* | NBRC | NBR95 |
| *mlk-1(ok2471)I* | CGC | RB1908 |

**Table S1 List of strains carrying extra chromosomal array used in this paper**

| Strain | Transgene | DNA Construct | Genetic Background | DNA concentration |
| --- | --- | --- | --- | --- |
| NBR95 | *shrEx472* | pNBRGWY124 (*Pser2prom4[4.1kb]::mScarlet*) | *N2* | 10ng/µl |
